# Entorhinal grid-like codes for visual space during memory formation

**DOI:** 10.1101/2024.09.27.615339

**Authors:** Luise P. Graichen, Magdalena S. Linder, Lars Keuter, Ole Jensen, Christian F. Doeller, Claus Lamm, Tobias Staudigl, Isabella C. Wagner

**Affiliations:** Department of Cognition, Emotion, and Methods in Psychology, Faculty of Psychology, University of Vienna, Vienna, Austria; University Medical Center Hamburg-Eppendorf (UKE), Hamburg, Germany; Centre for Human Brain Health, School of Psychology, University of Birmingham, Birmingham, United Kingdom; Max Planck Institute for Human Cognitive and Brain Sciences, Leipzig, Germany; Kavli Institute for Systems Neuroscience, Centre for Neural Computation, The Egil and Pauline Braathen and Fred Kavli Centre for Cortical Microcircuits, Jebsen Centre for Alzheimer’s Disease, Norwegian University of Science and Technology, Trondheim, Norway; Department of Psychology, Ludwig-Maximilians-Universität München, Munich, Germany; Donders Institute for Brain, Cognition, and Behaviour, Radboud University, Nijmegen, The Netherlands; Vienna Cognitive Science Hub, University of Vienna, Vienna, Austria; Centre for Microbiology and Environmental Systems Science, University of Vienna, Vienna, Austria

**Keywords:** Grid-like codes, entorhinal cortex, saccades, frontal eye fields, memory, functional magnetic resonance imaging (fMRI)

## Abstract

Eye movements, such as saccades, allow us to gather information about the environment and, in this way, can shape memory. In non-human primates, saccades are associated with the activity of grid cells in the entorhinal cortex. Grid cells are essential for spatial navigation, but whether saccade-based grid-like signals play a role in human memory formation is currently unclear. Here, human participants underwent functional magnetic resonance imaging (fMRI) and continuous eye gaze monitoring while studying scene images. Recognition memory was probed immediately thereafter. Results revealed saccade-based grid-like codes in the left entorhinal cortex while participants studied the scene images, a finding that was replicated with an independent data set reported here. The grid-related effects were time-locked to activation increases in the frontal eye fields. Most importantly, saccade-based grid-like codes were associated with recognition memory, such that grid-like codes were lower the better participants performed in subsequently recognizing the scene images. Collectively, our findings suggest an entorhinal map of visual space that is timed with neural activity in oculomotor regions, supporting memory formation.

## Introduction

Humans move their eyes to gather information about the environment. During natural viewing, eye movements, such as saccades, help to shift attention to the relevant features of a visual scene and can thereby shape memory formation (Bicanski & Burgess, 2019; Henderson, 2017; Lucas et al., 2019; Meister & Buffalo, 2016; Pertzov et al., 2009). Saccades are associated with the activity of grid cells in the entorhinal cortex, which are known to play a role in spatial navigation (Julian et al., 2018; Nau, Navarro Schröder, et al., 2018). Grid cells express multiple firing fields that are hexagonally arranged (Hafting et al., 2005). Killian and colleagues discovered visual grid cells in the entorhinal cortex of non-human primates that responded to multiple gaze positions (Killian et al., 2012) and saccade directions (Killian et al., 2015). In humans, it has been suggested that the firing properties of grid cell populations might relate to so-called “grid-like representations” or “grid-like codes”, which can be obtained non-invasively with functional magnetic resonance imaging (fMRI) and correspond to the strength of the hexadirectional modulation of the fMRI signal (Doeller et al., 2010; Kunz et al., 2019). Grid-like codes were linked to saccades during free visual search (Julian et al., 2018) and controlled visual tracking of a moving fixation target (Nau, Navarro Schröder, et al., 2018). Staudigl and colleagues employed magnetoencephalography (MEG) and reported a grid-like modulation of broadband high-frequency activity that was linked to saccades as individuals viewed scene images (Staudigl et al., 2018). Grid cells are thought to provide an internal spatial map of the current surroundings (Moser et al., 2008) that was shown to relate to spatial memory performance (Doeller et al., 2010; Kunz et al., 2015; Stangl et al., 2018; Wagner et al., 2023). Similarly, visual grid cells may provide us with a mental map to organize the spatial relationships of visually presented content and, in this way, guide memory formation (Killian & Buffalo, 2018). However, an effect of visual grid cells on memory has so far only been shown in non-human primates, where saccade-related grid cell activity revealed neural adaption (i.e., decreased activity) upon stimulus repetition (Killian et al., 2012, 2015), serving as an index for memory (Barron et al., 2016; Henson et al., 2000). This leaves open the question whether saccade-based grid-like codes in the human entorhinal cortex are relevant to memory formation.

The entorhinal cortex provides the major input into the hippocampus and is structurally connected to adjacent medial temporal areas (Garcia & Buffalo, 2020). In humans, it connects to the parahippocampal cortex, which is engaged in visual scene processing (Maass et al., 2015). In rodents and non-human primates, the entorhinal cortex receives input from primary and secondary visual areas (Campbell & Giocomo, 2018; Nadasdy et al., 2017). Brain regions associated with vision and oculomotion, such as the visual cortex and the frontal eye fields, play a central role in generating and coordinating saccades (Pierrot-Deseilligny et al., 2004; Prime et al., 2010). Recent studies revealed synchronized grid-like codes in human entorhinal and ventromedial prefrontal cortices that shared a similar grid orientation (Chen et al., 2021), as well as entorhinal-neocortical connectivity that was modulated by the magnitude of entorhinal grid-like codes (Wagner et al., 2023). These findings indicate that the entorhinal cortex might serve as a hub region, orchestrating information flow within entorhinal-neocortical networks (Buzsáki, 1996; Gerlei et al., 2021). However, how saccades and associated grid-like codes are integrated into this interregional dialogue remains to be elucidated.

To tackle these open questions, we examined data from two independent studies that were performed at different measurement sites (thus, yielding a “discovery” and a “validation” data set with *N* = 48 and *N* = 50, respectively). In both studies, human participants viewed scene images while we tracked their eye movements and measured fMRI. Immediately thereafter, participants were asked to discriminate previously studied from novel scenes during a recognition memory task (**Figure 1A**).

**Fig. 1.**
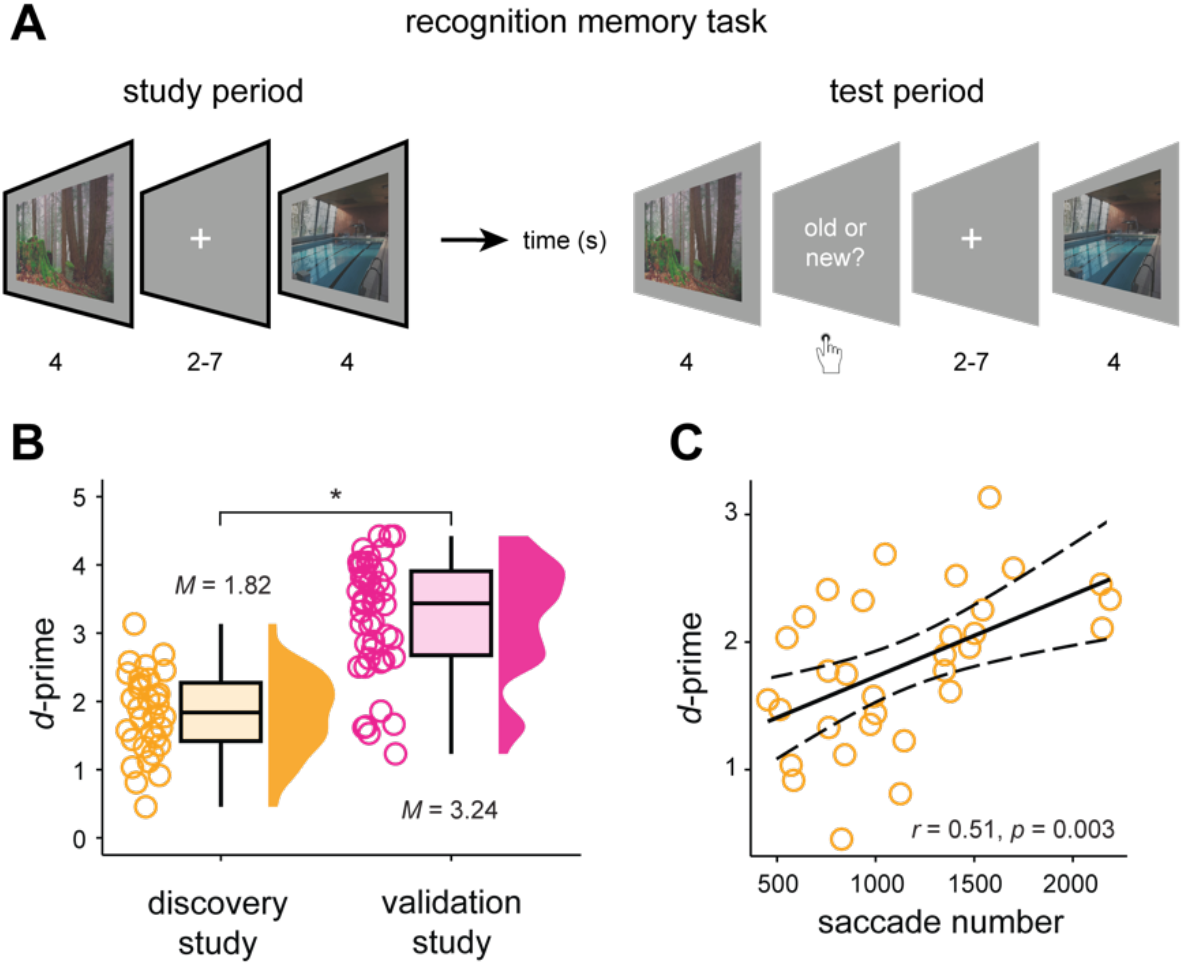
Recognition memory performance and saccades. A: Recognition memory task used in the discovery and validation study. Note that in the validation study, participants viewed each scene for 3 s (study period) and could provide their answer for 2 s (test period). The validation study also incorporated an additional test period that was performed in the behavioral lab after one week (delayed test, not depicted in the figure). B: Data points show the participant-specific *d*-prime values for both the discovery (orange) and the validation study (pink), and boxplots show the median (upper and lower borders mark the interquartile range, whiskers show minimum and maximum non-outlier values). C: The scatter plot shows the correlation between the total number of saccades per participant and *d*-prime for the discovery study (two-tailed, *N* = 32, *p* = 0.003; for the validation study, see Supplementary information, Figure S1B). The confidence interval (95% CI) is indicated by the dashed line. ^*^ = significant at *p* < 0.05.

We hypothesized that saccades during scene viewing should be coupled to grid-like codes in the entorhinal cortex and that the grid-related signals should be linked to individual variations in recognition memory performance. We further expected saccade-based grid-like codes in the entorhinal cortex to be tied to the activity in oculomotor regions that are known to be involved in saccade generation and coordination.

## Results

### Recognition memory performance and saccades

In both the discovery and validation study, participants completed a recognition memory task (**Figure 1A**) during which they studied scene images (study period) and were asked to discriminate previously viewed (“old”) from novel (“new”) scenes immediately after study (test period). Participants of the discovery study recognized more than two-thirds of the scene images correctly (∼ 70 % of 200 scenes, mean ± SEM, 140.91 ± 5.62) and performed, on average, 7.58 (± 0.40) saccades per scene. The average number of saccades during scenes that were later remembered (7.95 ± 0.40) was significantly higher than during scenes that were later forgotten (6.64 ± 0.43; paired-sample *t*-test, *N* = 32; *t*(31) = 8.262, Cohen’s *d* = 1.46, 95% confidence interval (CI) = [0.99, 1.63], *p*_*two-tailed*_ < 0.0001). In line with these results, we observed a positive correlation between individual recognition memory performance (*d*-prime, 1.82 ± 0.11; **Figure 1B, left panel**) and the number of saccades made when studying later remembered scenes (*r*_*Pearson*_ = 0.51, 95% CI = [0.20, 0.73], *p*_*two-tailed*_ = 0.003; **Figure 1C**). Thus, participants who used more saccades to explore later remembered scenes performed better in distinguishing old from novel material during the subsequent recognition memory test.

Similar to the discovery study, participants of the validation study recognized more than two-thirds of the scene images correctly (85% of 96 scenes, mean ± SEM, 81.78 ± 1.63) and performed, on average, 4.69 (± 0.14) saccades per scene. Again, the average number of saccades during scenes that were later remembered (4.74 ± 0.14) differed significantly from later forgotten scenes (4.31 ± 0.23; paired-sample *t*-test, *N* = 46; *p*_*two-tailed*_ = 0.008), and the number of saccades during later remembered scenes was significantly associated with individual recognition memory performance (*d*-prime, *r*_*Pearson*_ = 0.46, 95% CI = [0.20, 0.66], N = 46, *p*_*two-tailed*_ = 0.001; Supplementary Information, **Figure S1**). Overall, participants of the validation study showed high recognition memory performance (*d*-prime, 3.24 ± 0.13; **Figure 1B, right panel**), significantly better compared to the participants of the discovery study (discovery: *N* = 32, validation: *N* = 46; Wilcoxon rank sum test, *W* = 144, *p*_*two_tailed*_ < 0.001, *r* = -0.68). This is likely due to the fact that participants of the validation study viewed fewer scenes, were not faced with an additional (cover) task during encoding, and did not engage in a distractor task between the study and test periods (s. Methods).

### Saccades are associated with grid-like codes in the entorhinal cortex

Next, we tested whether saccades were linked to grid-like codes in the entorhinal cortex as participants viewed scenes during the study period. Based on previous findings in humans (Julian et al., 2018; Nau, Navarro Schröder, et al., 2018; Staudigl et al., 2018) and non-human primates (Killian et al., 2012), we expected significantly increased grid-like codes (i.e., testing for a 6-fold symmetrical modulation of the fMRI signal) in the entorhinal cortex time-locked to saccades. We thus turned towards the fMRI data and, for each participant, analyzed the saccades that occurred during scene viewing with respect to their directional angle. We then split the data of each individual into independent data halves (a) to estimate the individual, saccade-based grid orientation (i.e., the phase of the hexadirectional fMRI signal) in the first half of the data, and (b) to test the estimated grid orientation in the second half of the data (**Figure 2A**, and see Methods section for details). This allowed us to quantify the amount of saccade-based grid-like coding, predicting that a closer alignment between saccade direction and individual grid orientation would correspond to a stronger hexadirectional fMRI signal.

**Fig. 2.**
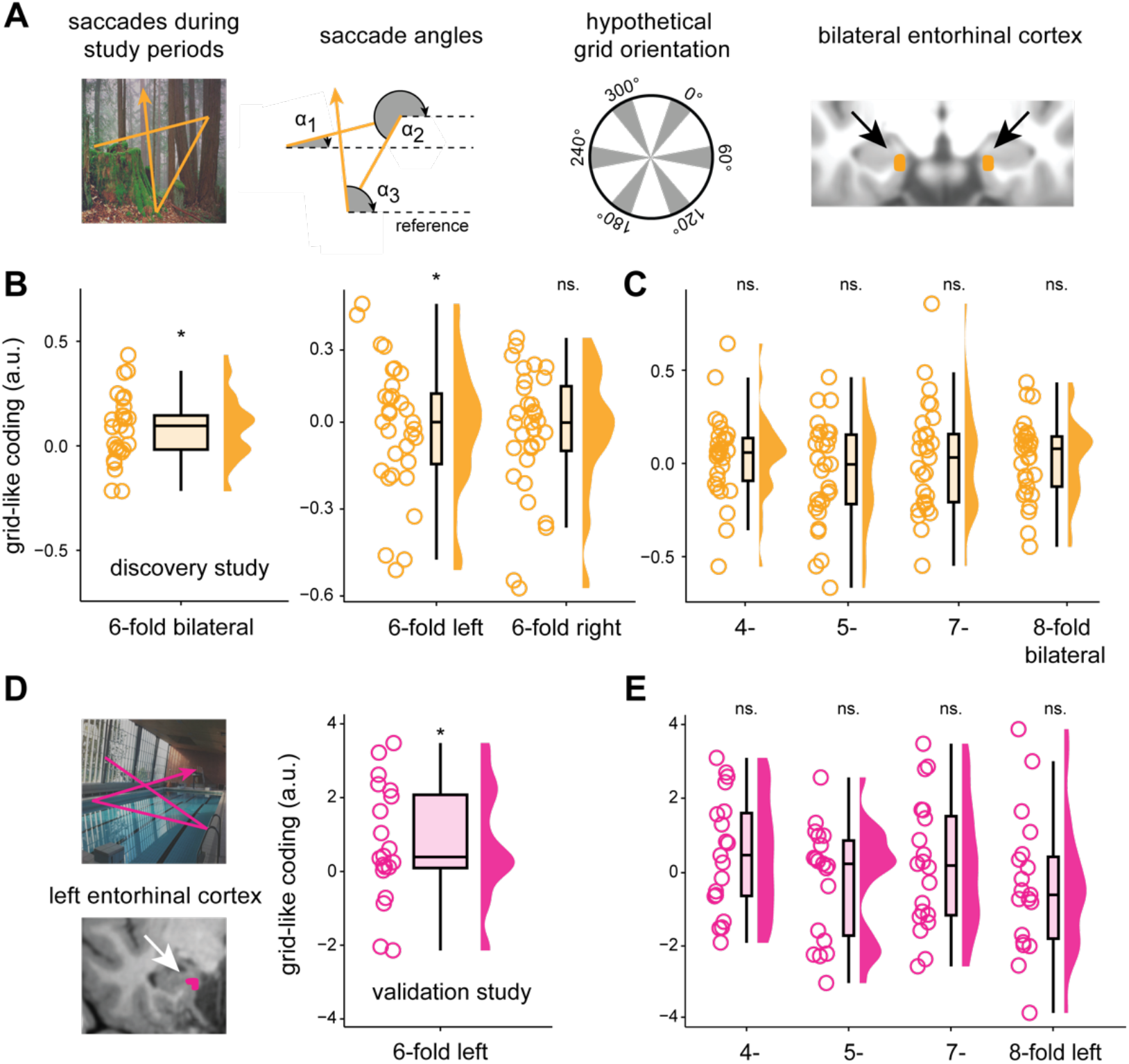
Saccades associated with grid-like codes in the entorhinal cortex. A: Discovery study (in orange), from left to right: Schematic trajectory of saccades during study period (exemplarily shown in orange and overlaid onto a scene image). Saccade directional angles referenced to an arbitrary point on the screen (dashed lines, angles α1, α2, α3 exemplarily shown in grey). Hypothetical grid orientation in 360° space with the main grid axes depicted in grey. Bilateral entorhinal cortex region-of-interest (ROI, in orange) projected onto the normalized T1-weighted structural image. We expected increased bilateral entorhinal cortex signal for saccades aligned to individual grid orientations. B, left panel: Magnitude of 6-fold grid-like codes in the bilateral entorhinal cortex (a.u., arbitrary units). B, right panel: Magnitude of 6-fold grid-like codes in left and right entorhinal cortices. C: Magnitude of grid-like codes for control symmetries (4-, 5-, 7-, and 8-fold periodicities) in the bilateral entorhinal cortex. D: Validation study (in pink), left panel: Schematic trajectory of saccades during study periods. Left entorhinal cortex ROI (in pink) superimposed on the T1-weighted structural image of one participant (note that grid-like codes for this study were analyzed in subject-native space). D, right panel: Magnitude of 6-fold grid-like codes in left entorhinal cortex. E: Magnitude of grid-like codes in left entorhinal cortex for control symmetries (4-, 5-, 7-, and 8-fold periodicities). Data points show individual grid-like codes during study periods, and boxplots show the median (upper and lower borders mark the interquartile range, whiskers show minimum and maximum non-outlier values). ^*^ = significant at *p* < 0.05; ns. = not significant.

In the discovery study data, we found significant grid-like coding in the bilateral entorhinal cortex, our predefined region-of-interest (ROI), for the 6-fold symmetrical model that was time-locked to saccades while participants studied the scene images (*N* = 29; bilateral entorhinal cortex: mean ± SEM, 0.084 ± 0.029, Wilcoxon test, *V* = 333.5, *p*_*one-tailed*_ = 0.0063, Cohen’s *d* = 0.53; **Figure 2B, left panel**). If this effect was actually related to grid-like codes, we reasoned that it should not be present for different symmetrical models (testing for a 4-, 5-, 7-, or 8-fold symmetry). Indeed, when we repeated the analysis with these different symmetries, results showed no evidence for significantly increased grid-like codes of these types in the entorhinal cortex (**Figure 2C**). We also tested whether grid-like codes were present in other areas known to be involved in memory and visuo-oculomotor processing (such as the hippocampus, anterior thalamus, frontal eye fields, and visual cortex). There were no significantly increased grid-like codes in any of these regions (Supplementary information, **Figure S2**), corroborating that our findings are specific to the entorhinal cortex in which grid cells were previously reported (Doeller et al., 2010). Additional follow-up analysis showed that the effect was more prominent for grid-like codes in the left than in the right entorhinal cortex (*N* = 29; left entorhinal cortex: mean ± SEM, 0.068 ± 0.034, Wilcoxon test, *V* = 300, *p*_*one-tailed*_ = 0.038, *d* = 0.37; right entorhinal cortex: 0.051 ± 0.031, Wilcoxon test, *V* = 275, *p*_*one-tailed*_ = 0.052, *d* = 0.30; **Figure 2B, right panel**).

We performed additional control analyses to rule out alternative explanations for our findings. First, we repeated the main analysis but reversed the estimation and test data halves (thus, we estimated individual grid orientations on the second data half and tested grid-like codes in the first data half). This gave virtually identical results, highlighting that our findings were independent of the specific data partitioning scheme (*N* = 29; bilateral entorhinal cortex: mean ± SEM, 0.087 ± 0.023, Wilcoxon test, *V* = 340, *p*_*one-tailed*_ = 0.0041, *d* = 0.45; Supplementary information, **Figure S3A**). Second, to show that grid-like codes were not linked to the number of saccades that participants made, we correlated the average number of saccades across all trials with the individual magnitude of entorhinal grid-like coding. Results confirmed that the magnitude of grid-like codes was not related to saccade numbers (*N* = 29; bilateral entorhinal cortex: *p*_two-tailed_ = 0.386). Third, the estimation of grid orientations can be biased by the saccade durations (in ms) along the different saccade directions on the computer screen (in other words, it is not possible to estimate individual grid orientations for a specific saccade direction if the participant never made saccades in that direction). We found that there was a significant bias in saccade durations across directional bins in 8 out of 29 participants (all *p* < 0.05). For this subset of participants, we then randomly excluded 10% of saccades in an iterative process until we could ensure an even distribution of saccade durations across the different saccade directions (all *p* > 0.05), and repeated the main analysis. Once again, this confirmed our result of increased saccade-based grid-like codes in the entorhinal cortex (Supplementary information, **Figure S3B**). Fourth, we checked whether significantly increased grid-like codes were tied to an overall increase in the entorhinal cortex BOLD signal that could reflect individual differences in the signal-to-noise ratio of the fMRI data (thus, answering the question of whether stronger grid-like codes would be linked to a higher overall BOLD signal in the entorhinal cortex), but this was not the case (*p* = 0.083, Supplementary information, **Figure S4**).

Building on the abovementioned findings from the discovery study, we then leveraged the data from the validation study to test for saccade-based grid-like codes in this independent participant sample. Since grid-like codes were more prominent in the left entorhinal cortex (**Figure 2B, right panel**), we focused the analysis on the left and right entorhinal cortices separately (we only performed this analysis on a subset of 20 participants for which the entorhinal cortex masks comprised at least 14 voxels to match the entorhinal cortex mask size of the discovery study, see Methods section for details, and as we have done previously; Wagner et al., 2023). Once again, we detected significant saccade-based grid-like codes in the left but not in the right entorhinal cortex while participants studied scene images (*N* = 20; left entorhinal cortex: mean ± SEM, 0.746 ± 0.353, Wilcoxon test, *V* = 162, *p*_*one-tailed*_ = 0.01638, *d* = 0.472; right entorhinal cortex: mean ± SEM, -0.122 ± 0.393, Wilcoxon test, *V* = 94, *p*_one-tailed_ = 0.663, *d* = -0.07, *r* = 0.092; **Figure 2D**). Grid-like codes were not significant for any of the control symmetries (*N* = 20; left entorhinal cortex; 4-fold: mean ± SEM, -0.049 ± 0.634; 5-fold: -0.038 ± 0.387; 7-fold: 0.169 ± 0.390; 8-fold: -0.702 ± 0.525; Wilcoxon test, all *p*_*one-tailed*_ > 0.05; **Figure 2E**). In summary, across two independent data sets, we found significantly increased saccade-based grid-like codes in the left entorhinal cortex while participants studied scene images.

### Saccade-based grid-like codes in the entorhinal cortex are lower at better recognition memory

Our main goal was to clarify the role of saccade-based grid-like codes in memory formation. Previous studies reported mixed results regarding the relationship between entorhinal grid-like codes and behavior. In short, increased grid-like codes were associated with better spatial navigation performance of human participants in an object-location memory task (i.e., lower drop error when placing an object at its correct location; Doeller et al., 2010; Kunz et al., 2015; Stangl et al., 2018). When observing a virtual demonstrator, increased grid-like codes were associated with lower navigation performance (Wagner et al., 2023), and a similar relationship was found for directional coding in the human medial temporal lobe (Nau et al., 2020). Saccade-based grid-like codes were shown to be positively associated with self-reported navigation ability (Julian et al., 2018). In non-human primates, only some of the detected visual grid cells showed neural adaption (i.e., decreased activation) upon stimulus repetition (Killian et al., 2012; Meister & Buffalo, 2016), leaving it open whether saccade-based entorhinal grid-like codes are related to human memory formation as well.

Using data from the discovery study, we tested whether individual variations in the magnitude of saccade-based grid-like codes during study periods would scale with individual differences in recognition memory performance (as indexed by *d*-prime). Results showed a significantly negative association between grid-like codes and *d*-prime values (*N* = 29, *r*_*Pearson*_ = -0.51, 95% CI = [-0.740, -0.182], *p*_*two-tailed*_ = 0.004; **Figure 3A, left panel**). In other words, lower saccade-based grid-like codes were correlated with better recognition memory performance across participants.

**Fig. 3.**
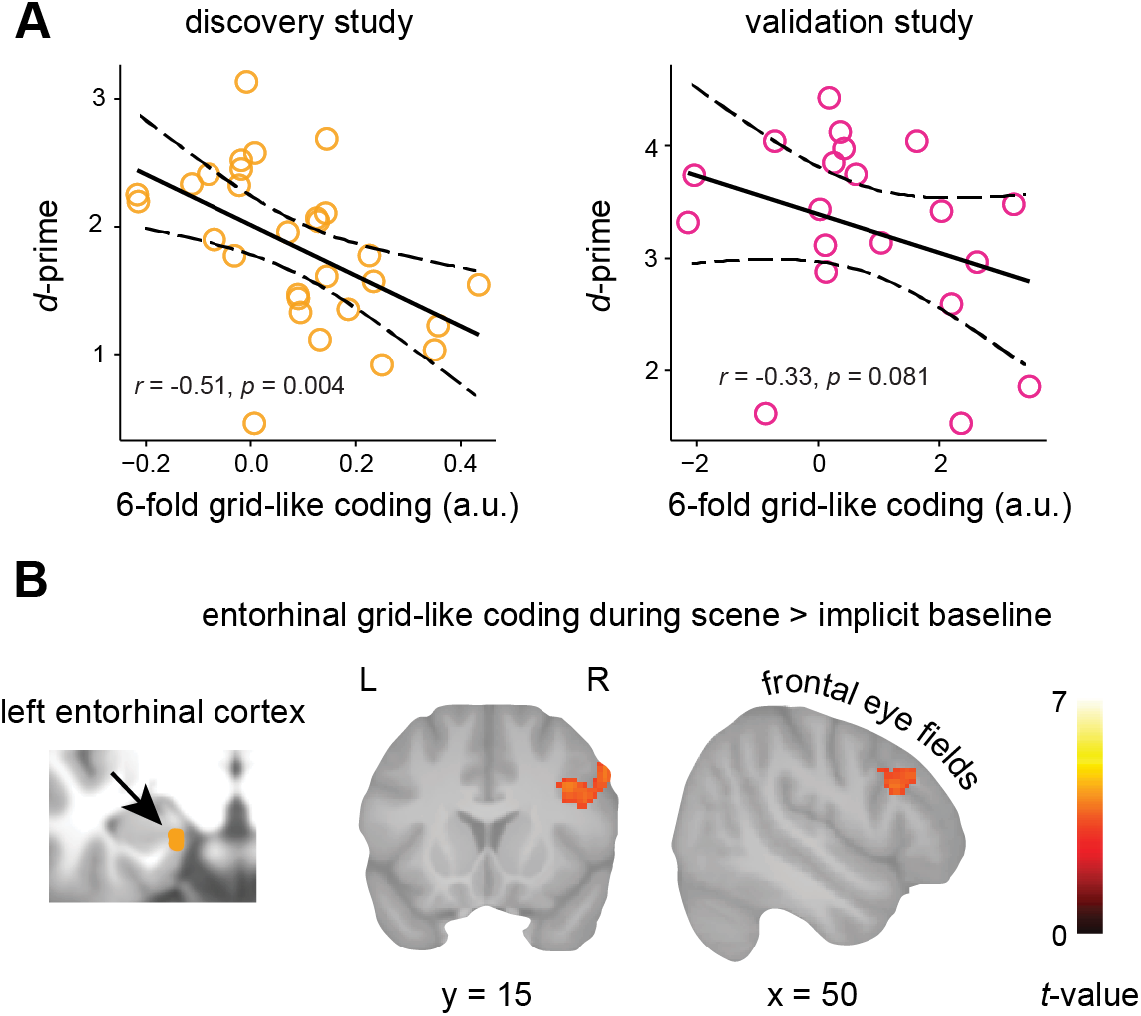
Saccade-based grid-like codes are lower at better recognition memory performance and are time-locked to frontal eye field activation. A: The scatter plots display the relationship between the magnitude of grid-like codes (a.u., arbitrary units) and individual recognition memory performance (*d*-prime) for both the discovery study (in orange, two-tailed, *N* = 29, *p* = 0.004, 95% confidence interval CI indicated by the dashed line) and the validation study (in pink, one-tailed, *N* = 20, *p* = 0.081, 95% CI indicated by the dashed line). ^*^ = significant at *p* < 0.5. B: In the discovery study, neural activity in the frontal eye fields co-varied with the magnitude of saccade-based grid-like codes in the entorhinal cortex. Results are shown at *p* < 0.05 FWE-corrected at cluster-level (cluster-defining threshold of *p* < 0.001, cluster extent = 116 voxels).

Several control analyses confirmed this brain-behavior relationship. First, we could show that the result did not stem from specific saccade patterns. In other words, it could have been the case that participants with better recognition memory (thus, higher *d*-prime) made shorter saccades, which could have caused lower grid-like codes, but this was not the case (*p* = 0.717; Supplementary information, **Figure S5A**). Second, we could show that the result was not driven by differences in entorhinal BOLD activation. It could have been the case that participants with higher *d*-prime showed lower BOLD changes during the study period, as better memory can be associated with activation decreases in the involved areas (Wagner et al., 2021), causing lower grid-like codes. However, this was also not the case (*p* = 0.094; Supplementary information, **Figure S5B**).

We repeated the same analysis with data from the validation study, for which recognition memory performance was measured twice, immediately after the study period (immediate test) and one week later (delayed test). Results revealed a tendency towards lower grid-like codes in the left entorhinal cortex at better recognition memory performance across participants at the immediate test, but this was not significant (*d*-prime; *N* = 20, *r*_*Pearson*_ = -0.33, 95% CI = [-1, 0.061], *p*_*one-tailed*_ = 0.081; **Figure 3A, right panel**). A similar picture emerged when taking into account the performance in the delayed recognition memory test (*d*-prime; *N* = 20, *r*_*Pearson*_ = -0.28, 95% CI = [-1, 0.108], *p*_*one-tailed*_ = 0.114). Once again, we suspect that this might be due to differences in task design between the discovery and validation studies (in the latter, participants studied fewer scenes and were not required to solve an additional task between the study and test periods). While the effect was only robust in the discovery study, analyses suggest a potential association between lower saccade-based grid-like codes and successful memory formation.

### Saccade-based entorhinal grid-like codes are time-locked to neural activity in the frontal eye fields

Saccade generation and coordination are associated with neural activity in a set of brain regions that appear engaged in visual processing and oculomotion, including the visual cortex and the frontal eye fields (Johnston & Everling, 2008; Munoz & Everling, 2004). Consequently, we reasoned that saccade-based entorhinal grid-like codes would be coupled to activation changes in this wider network of brain regions. To reveal whether BOLD activation varied as a function of saccade-based entorhinal grid-like codes, we adopted a whole-brain approach. We examined the data from the discovery study, averaging the magnitude of entorhinal grid-like coding across all saccades within a single scene trial. We then modeled all scene trials during the study period and included the magnitude of trial-wise grid-like coding as a parametric modulator in a group-based analysis (see Methods section).

Results from the whole-brain analysis showed that, as participants viewed scene images, larger saccade-based grid-like codes were coupled to increased activation specifically in the left frontal eye fields. In other words, if participants viewed scene images and made saccades that were aligned with their individual entorhinal grid orientation (i.e., resulting in larger grid-like codes for that trial), activation in the left frontal eye fields was high as well (*p* < 0.05 FWE-corrected at cluster level using a cluster-defining threshold of *p* < 0.001, cluster size = 116 voxels; *x* = 55, *y* = 22, *z* = 31; **Figure 3B**).

## Discussion

In the present study, we investigated whether saccade-based grid-like codes in the human entorhinal cortex play a role in human memory formation. Leveraging data from two independent data sets, we consistently identified saccade-based grid-like codes in the left entorhinal cortex while participants studied scene images. Interestingly, in the discovery study, grid signals appeared lower at better individual recognition memory and higher grid signals were associated with increased neural activation in the frontal eye fields. Together, these findings are the first to highlight saccade-based entorhinal grid-like codes as a potential player in human memory formation and reveal their link to the frontal eye fields that are crucial for eye movement coordination.

We hypothesized that saccades during scene viewing were coupled to grid-like codes in the entorhinal cortex and that the grid-related signals would be linked to individual variations in recognition memory performance. In line with this prediction, we found significantly increased entorhinal grid-like codes related to saccades in two independent studies. This is consistent with reports of grid cells in the entorhinal cortex of non-human primates that were shown to respond to multiple gaze locations and saccade directions during free visual exploration (Killian et al., 2012, 2015). Similarly, grid-like codes in the human entorhinal cortex were discovered while individuals performed a visual search task (Julian et al., 2018), a visual tracking task (Nau, Navarro Schröder, et al., 2018), or while freely viewing scene images (Staudigl et al., 2018). By encoding saccade directions, grid-like signals may represent relational information between the different elements of a visual scene (Bicanski & Burgess, 2019), potentially providing us with a mental framework for the organization of visual memory reminiscent of the cognitive map for physical space (Killian & Buffalo, 2018; Nau, Julian, et al., 2018). Mapping visual space to guide navigation should be highly adaptive for all animals that use vision as their primary sense for exploration, including humans.

Crucially, we found that saccade-based grid-like codes were lower the better participants were able to correctly discriminate old from novel scene images in a subsequent recognition memory task (we also detected a similar trend in the validation data set, albeit not significant, which is likely due to the fact that these participants encoded substantially fewer scene images). To the best of our knowledge, only one previous paper described a relationship between visual grid cell activity and visual memory in non-human primates (Killian et al., 2012). The authors showed that grid cells located in the anterior entorhinal cortex displayed neural adaption upon stimulus repetition (i.e., decreased activation as a surrogate marker of memory; Barron et al., 2016; Henson et al., 2000). Other work on the relationship between (non-visual) grid-like codes and behavior yielded mixed results: Some studies have linked lower (Wagner et al., 2023) or higher grid-like codes (Doeller et al., 2010; Kunz et al., 2015; Stangl et al., 2018) to better spatial memory performance, while others reported no significant brain-behavior relationship (Horner et al., 2016; Nau, Navarro Schröder, et al., 2018). A possible explanation for the negative brain-behavior result is that participants who were better at recognizing the scene images may have used a different strategy to complete the task. Rather than relying only on visuospatial encoding, they may have drawn upon prior knowledge. For instance, when viewing a specific beach scene, they might have recalled a recent beach vacation with highly similar scene features. Such integration of prior knowledge, or memory “schemas”, could have facilitated scene encoding and thus enhanced participant’s recognition memory (Bonasia et al., 2018; Maguire et al., 1999; Van Kesteren et al., 2012). In turn, the processing of spatial relations between individual scene features, intrinsic to visuospatial encoding, may have been less effective, potentially manifesting in reduced grid-like coding for visual space (Bonasia et al., 2018; Van Kesteren et al., 2013). Another possible reason is that individuals with good recognition memory might display less neural activation during memory formation, thus, encoding “more efficiently” (Neubauer & Fink, 2009). For instance, increased memory performance due to memory training was associated with reduced neural activation in brain areas relevant to mental navigation, including the posterior hippocampus, and retrosplenial cortex (Wagner et al., 2021). In macaques, long-term practice was associated with reduced glucose uptake while maintaining neural activity levels (Picard et al., 2013). We previously showed that individuals who exhibited less entorhinal grid-like coding and entorhinal-cortical connectivity when observing a demonstrator moving through virtual space performed better when later retracing the demonstrator’s path (Wagner et al., 2023). Nau and colleagues found that fMRI-based directional coding in the medial temporal lobe was weaker in participants with better memory performance in a virtual spatial navigation task as quantified by a smaller drop-error when trying to replace objects to their locations (Nau et al., 2020). Note, however, that the correlation between grid-like coding and overall BOLD signal was not significant in our data set (Supplementary information, **Figure S4**). To test whether the negative brain-behavior relationship was caused by potentially confounding factors, such as specific saccade patterns (e.g., shorter saccade durations) or reduced activity profiles (i.e., neural adaption) in individuals with better recognition memory, we performed several control analyses and were able to rule out these alternative explanations. Consequently, our findings reinforce the notion that grid-like codes, which are typically discussed in the context of spatial navigation, also play a role in the representation of visual space and contribute to memory formation.

We next hypothesized that saccade-based grid-like codes in the entorhinal cortex were tied to neural activity in visuo-oculomotor regions that are known to be involved in saccade generation and coordination, such as the visual cortex or frontal eye fields (Prime et al., 2010). Indeed, we observed that increased grid-like coding in the entorhinal cortex was related to an increased activation in the frontal eye fields. In other words, if participants made saccades that were aligned with their individual grid orientation, activation in the frontal eye fields was high as well. The entorhinal cortex and the frontal eye fields are closely connected, and are embedded in a set of regions spanning from visual (Nadasdy et al., 2017) to medial temporal areas (Burwell & Amaral, 1998; Van Strien et al., 2009). Disynaptic pathways between these areas provide the ideal infrastructure to interface between regions for memory and oculomotion (Ryan & Shen, 2020). Ryan and colleagues (2020) modeled the functional dynamics between these regions and demonstrated that medial temporal lobe lesions disrupted signal transmission to different oculomotor areas, including the frontal eye fields (Ryan et al., 2020). This is in line with findings from patients with Alzheimer’s Disease (AD) who show the first signs of neurodegenerative changes in the entorhinal cortex (Braak & Del Tredici, 2015). Importantly, AD patients display altered eye movement patterns, with saccades lacking accuracy and speed when directed towards, or when suppressing movement to, a predefined target (Molitor et al., 2015; Opwonya et al., 2022, 2023; Readman et al., 2021). This suggests that the entorhinal cortex provides relevant visuospatial input into the frontal eye fields, thereby informing the planning and execution of subsequent saccades. In turn, saccades shape the activity in the medial temporal lobe (Jutras et al., 2013; Liu et al., 2017). Dynamic causal modeling revealed that free viewing during mental scene construction (as opposed to a restricted viewing condition) enhanced excitatory functional connections from the medial temporal lobe to the frontal eye fields, indicating the influence of saccadic activity on the interaction between these regions (Ladyka-Wojcik et al., 2022). Furthermore, transcranial magnetic stimulation of the frontal eye fields was shown to disrupt spatial working memory (Prime et al., 2010). Considering these findings, saccade-based entorhinal grid-like codes may not only encode the spatial relationships between visual features but also interact with the frontal eye fields to guide the computation of future saccade directions and to support perception and memory (Bicanski & Burgess, 2019). In doing so, grid signals may help coordinate the interplay between medial temporal and visuo-oculomotor regions in support of memory formation.

Even though grid-like codes have been consistently reported in prior and our present work, there is an active debate about the extent to which fMRI-based grid-like codes reflect the underlying cellular activity (Kunz et al., 2019). Single-cell recordings in humans have identified grid cells in the entorhinal cortex during virtual navigation (Jacobs et al., 2013), and entorhinal grid-like codes have been linked to saccadic eye movements (Julian et al., 2018; Nau, Navarro Schröder, et al., 2018). MEG and intracranial electroencephalography (EEG) recordings revealed grid-like modulation of visual space in the human anterior medial temporal lobe (Staudigl et al., 2018). However, direct evidence connecting grid cell recordings in humans to saccades during visual exploration is currently missing. Bicanski and Burgess proposed a computational model in which grid cells encode trajectories between salient stimulus features of a visual scene, offering a possible explanation for their role in memory encoding (Bicanski & Burgess, 2019). To gain valuable insights into their function in visual processing and memory encoding, grid-like codes should be explored across different stimulus types and various levels of complexity, beyond scene images. Another challenge is posed by the generally slower temporal resolution of the fMRI-based BOLD signal, rendering it unlikely that measured fluctuations in brain activity can be tied to single saccades (i.e., fast-occurring events with millisecond-duration). While we employed a short repetition time (TR) to maximize the temporal resolution in the discovery study (657 ms), measurements in the validation study were based on a longer TR (2029 ms). This may account for the generally weaker effects observed in that data set (i.e., one TR likely included several saccades, potentially yielding lower grid-like codes due to averaging across multiple saccades that were (mis-)aligned with the individual grid orientation). Nevertheless, our results are backed up by previous studies that assessed saccade-based grid-like codes with fMRI (Julian et al., 2018; Nau, Navarro Schröder, et al., 2018), as well as by our numerous control analyses that corroborate the validity and stability of the findings.

To conclude, we identified grid-like codes in the entorhinal cortex that were locked to saccades. Across individuals in the discovery study, these saccade-based grid-like codes during scene viewing were lower at better subsequent recognition memory, suggesting that grid signals contributed to memory formation. Moreover, the magnitude of grid-like coding was coupled to increases in neural activation of the frontal eye fields, a brain region that is known to be involved in saccade generation and coordination. Our findings are the first to show that saccade-based grid-like codes in the entorhinal cortex play a role in human memory formation, highlighting interregional coordination of neural activity that is time-locked to the internal map of visual space.

## Materials and methods

### Data set obtained at the Donders Institute – the “discovery study”

#### Study setup

This study belonged to a larger project examining the effects of eye movements on memory processing (performed at the Donders Institute for Brain, Cognition and Behaviour, Nijmegen, The Netherlands). In two separate sessions, participants underwent MEG (not reported here, but see Staudigl et al., 2017) and fMRI (see also Wagner et al., 2022) while monitoring their eye movements, and while performing a recognition memory task. The order of fMRI/MEG sessions was balanced across participants and involved parallel task versions to avoid training effects.

#### Participant sample

Forty-eight participants volunteered for this study. Sixteen participants were excluded due to not completing the study (7 individuals), excessive motion (4 individuals), technical problems during the data recording (3 individuals), a low number of identified saccades during the fMRI session (1 individual, < 30 detected saccades per condition), or low recognition memory performance during the MRI session (1 individual, false alarms > correct rejections). The final sample thus comprised 32 participants (23 females, age range 18-30 years, mean age = 23 years, 32 right-handed). All individuals were healthy and did not report any history of neurological and/or psychiatric disorders, had normal or corrected-to-normal vision, and provided written informed consent before the start of the experiment. The study was reviewed and approved by the local ethics committee (Commissie Mensgebonden Onderzoek, region Arnhem-Nijmegen, The Netherlands; reference number CMO-2014/288).

#### Recognition memory task

During the study period, participants were instructed to memorize 200 scene images (100 indoor, 100 outdoor). Images were resized to a dimension of 1024 × 768 pixels and were presented on a black background. Each scene was shown for 4 s during which participants could freely view the image. To ensure attention to each scene, participants were asked to judge whether the image depicted an indoor or outdoor scenario via button press during the subsequent fixation period (2125, 4125, or 7125 ms, 80/80/40 distribution across the 200 trials, pseudo-randomized), after which the next scene appeared. The order of scenes was pseudorandomized, with no more than four scenes of the same type (indoor/outdoor) shown consecutively. The study period was followed by a distractor task (i.e., solving simple mathematical problems, 1 min) and a rest period (3 min).

During the test period, participants viewed all scene images that were shown during the previous study period, intermixed with 100 novel scene images (half of them indoor/outdoor; i.e., a total of 300 scene images were presented). The assignment of scene images to study or test periods was counterbalanced across participants. Scenes were presented for 4 s each and were followed by a 6-point rating scale that required participants to indicate whether they recognized the scene as “old” or “new” (self-paced; the scale ranged from (1) “very sure old” to (6) “very sure new”), and a fixation period until the next trial started (2125, 4125, or 7125 ms, 80/80/40 distribution across the 300 trials, pseudo-randomized). The test period was divided into 2 blocks separated by a short break. After completing the task, participants were asked to fixate on different locations on the screen to evaluate eye tracker accuracy (5 min), followed by the structural scan.

#### Recognition memory performance (d-prime)

Trials were grouped into four bins based on individual performance during the recognition memory test: (1) scenes that were correctly judged as “old” (i.e., hits, collapsing across confidence ratings 1-3, mean ± standard error of the mean (SEM): 140.9 ± 5.6 trials); (2) scenes that were correctly judged as “new (i.e., correct rejections, collapsing across confidence ratings 4-6, 85.8 ± 1.8 trials); (3) scenes that were incorrectly judged as “old” (i.e., false alarms, collapsing across confidence ratings 1-3, 14.2 ± 1.8 trials); (4) scenes that were incorrectly judged as “new” (i.e., misses, collapsing across confidence ratings 4-6, 59.1 ± 5.6 trials). None of the participants displayed any actually missed trials without button presses. Individual hit and false alarm rates were *z*-scored, and recognition memory performance (*d*-prime) was calculated as [*z*(hits) – *z*(false alarms)].

#### Eye tracking data acquisition, analysis, and saccade detection

To capture saccadic eye movements, we recorded horizontal and vertical eye gaze and pupil size, using a video-based infrared eye tracker (EyeLink 1000 Plus, SR Research, Ontario, Canada). Before recording, raw eye movement data was mapped onto screen coordinates by means of a calibration procedure. Participants sequentially fixated on nine fixation points on the screen, arranged in a 3 × 3 grid. This was followed by a validation procedure during which the nine fixation points were presented once more while the differences between the current and previously obtained gaze fixations (from the calibration period) were measured. The calibration settings were accepted if these differences were < 1° of visual angle, and the eye tracker recording was started.

Eye tracking data was processed using Fieldtrip (https://www.fieldtriptoolbox.org). Saccadic eye movements were identified by transforming vertical and horizontal eye movements into velocities, whereby velocities exceeding a threshold of 6 x the standard deviation (*SD*) of the velocity distribution and with a duration of > 12 ms were defined as saccades (Engbert & Kliegl, 2003). Saccade onsets during trials of the study period (i.e., during the presentation of scene images) were defined as events-of-interest. Only saccades that followed a minimum fixation period of 25 ms were included. Saccades that were followed or preceded by blinks (+/-100 ms) were excluded (blinks were defined as large deflections in pupil diameter: mean ± 5 standard deviations; eye tracking data in the vicinity of blinks is unreliable due to saturation effects). Trials with more than 25% of missing eye tracker data were discarded. We detected a total of 48510 saccades in the eye tracking data (*N* = 32; average number of saccades per participant, mean ± SEM: 1515.94 ± 80.95 saccades).

#### MRI data acquisition

Imaging data were collected at the Donders Institute for Brain, Cognition and Behaviour (Nijmegen, The Netherlands), using a 3T Prisma Fit scanner (Siemens, Erlangen, Germany) equipped with a 32-channel head coil. We acquired on average 2456 (± 5.3) T2*-weighted blood oxygen level-dependent (BOLD) images during the study period of the recognition memory task, using the following echo-planar imaging (EPI) sequence: repetition time (TR) = 657 ms, echo time (TE) = 30.8 ms, multi-band acceleration factor = 8, 72 axial slices, interleaved acquisition, field of view (FoV) = 174 × 174 mm, 72 × 72 matrix, flip angle = 53°, slice thickness = 2.4 mm, no slice gap, voxel size = 2.4 mm isotropic. The structural image was acquired using a standard magnetization-prepared rapid gradient-echo (MPRAGE) sequence with the following parameters: TR = 2300 ms, TE = 3.03 ms, FoV = 256 × 256 mm, flip angle = 8°, voxel size = 1 mm isotropic.

#### MRI data preprocessing

The fMRI data were processed with SPM8 in combination with MATLAB (The Mathworks, Natick, MA, USA). The first 12 volumes were excluded to allow for T1-equilibration. The remaining volumes (of both the study and test periods) were realigned to the mean image. The structural scan was co-registered to the mean functional image and was segmented into grey matter, white matter, and cerebrospinal fluid using the “New Segmentation” algorithm. All images (functional and structural) were then spatially normalized to the Montreal Neurological Institute (MNI) EPI template using Diffeomorphic Anatomical Registration Through Exponentiated Lie Algebra (DARTEL; Ashburner, 2007a), and functional images were further smoothed with a 3D Gaussian kernel (6 mm full-width at half-maximum, FWHM).

#### Region-of-interest (ROI) definition

For the analysis of grid-like codes, left and right posterior medial entorhinal cortex masks were based on Maass et al., 2015. Masks were binarized and co-registered to the mean functional image of one participant (Maass, bilateral entorhinal cortex: 25 voxels, left entorhinal cortex: 14 voxels, right entorhinal cortex: 18 voxels). To validate the quality of the co-registration, the overlap between each mask and the corresponding (co-registered) structural and mean functional image was visually assessed for each participant.

To test for potential grid-like codes in control regions, we defined additional ROIs that are known to be involved in memory, visuo-spatial processing, and oculomotor control, but for which no significant grid-like codes have been detected. This included the hippocampus, anterior thalamus, frontal eye fields, and visual cortex. The hippocampus and visual cortex were defined based on bilateral anatomical masks of the Automatic Anatomical Labeling (AAL) atlas (Tzourio-Mazoyer et al., 2002; hippocampus = 1148 voxels, visual cortex = 1860 voxels). To delineate the anterior thalamus, we used the stereotactic mean anatomical atlas provided by Krauth and colleagues (Krauth et al., 2010; © University of Zurich and ETH Zurich, Axel Krauth, Rémi Blanc, Alejandra Poveda, Daniel Jeanmonod, Anne Morel, Gábor Székely), which is based on histological, cytoarchitectural features defined *ex vivo* (Morel, 2007). We specified the anterior thalamus by combining the bilateral anterior dorsal, -medial, and -ventral nucleus masks (59 voxels). The frontal eye fields were defined by contrasting memory-related activity during scene encoding across all participants (later remembered > later forgotten). The resulting cluster peak coordinate (*x* = 43, *y* = 7, *z* = 29) was surrounded by a 10 mm sphere and was mirrored to create a bilateral ROI (320 voxels). See Supplementary information, **Figure S2A**).

#### Analysis of grid-like codes

Grid-like codes were analyzed using the openly available Grid Code Analysis Toolbox (GridCAT, software version 1.0.4, https://www.nitrc.org/projects/gridcat; (Stangl et al., 2017) which is based on the procedures developed by Doeller et al. (2010).

Saccades during the study period of the recognition memory task were defined as events-of-interest. We then leveraged the General Linear Model (GLM) to model the BOLD response time-locked to saccade onsets. All saccades were estimated with stick functions (duration = 0 seconds) and were convolved with the SPM default canonical hemodynamic response function (HRF). To account for noise due to head movement, we included the six realignment parameters, their first derivatives, and the squared first derivatives into the design matrix. A high-pass filter with a cutoff at 128 s was applied.

Analysis of grid-like codes progressed in two steps pertaining to estimating and testing individual grid orientations. We partitioned the data into two equally-sized data halves (i.e., corresponding to two separate regressors that contained the saccades of the estimation or test data sets, respectively). During step 1 (GLM 1), saccade-related activity of the estimation data set (i.e., the first regressor) was modulated by the respective saccade direction. This was calculated as saccade angle (α_t_) relative to a predefined reference point and was modeled using two parametric modulators, sin(α_t_*6) and cos(α_t_*6), that converted directional information into 60° space, reflecting the hypothesized 6-fold rotational symmetry in the fMRI signal (presumably due to the firing patterns of underlying grid cells). The voxel-wise beta estimates, β_1_ and β_2_, of the two parametric modulators were then extracted, and the mean grid orientation within the respective ROI was calculated using arctan[mean(β_1_)/mean(β_2_)]/6 (i.e., converting directional information back into 360° space).

During step 2, the estimated grid orientations were then tested in a second GLM (GLM 2) that was virtually identical to the abovementioned model, but with the exception that the saccades within the first regressor (i.e., the estimation data set) were unmodulated, while the saccades in the second regressor (i.e., the test data set) were parametrically modulated by the difference between the respective saccade angle (α_t_) and the individual ROI-based grid orientation (φ) using cos[6*(α_t_-φ)]. In other words, a smaller difference between α_t_ and φ should result in increased grid-like codes since the saccade direction is aligned with the individual grid orientation. The beta values from the parametric modulator were then extracted for all voxels within the ROI and were averaged to produce the mean amount of grid-like coding.

ROI-based grid-like code data were analyzed using a set of Wilcoxon-tests in R (software version 4.3.0; https://www.r-project.org; R stats version 3.6.2). We hypothesized that significant grid-like coding in the entorhinal cortex should be associated with a 6-fold rotational symmetry of the fMRI signal in the entorhinal cortex. This is based on the assumption that participants cross more grid cell firing fields as they perform saccades aligned with the underlying grid axes. The choice of the statistical test thus reflected an *a priori* expectation, which is why we adopted an α-level of 0.05 (one-tailed). Effect sizes were calculated as Cohen’s *d*. Additionally, we applied Bonferroni-correction to account for multiple comparisons (1 entorhinal cortex ROI and 4 control ROIs), using a threshold of α_Bonferroni_ = 0.05/5 ROIs = 0.01 or (6-fold symmetry and 4 control symmetries), using a threshold of α_Bonferroni_ = 0.05/5 ROIs = 0.01, respectively. Grid-like coding values exceeding the median value ± 3 × the median absolute deviation (MAD) were excluded from the analyses. We chose this method because the mean and standard deviation are particularly sensitive to outliers whereas the median is not (Leys et al., 2013).

#### Whole-brain activation modulated by saccade-based entorhinal grid-like codes

We performed additional analyses to test whether the activity of entorhinal grid-like codes modulated voxel-wise changes in whole-brain activation. The amount of grid-like coding of each saccade was taken from the results of GLM 2 (this GLM had tested the previously estimated grid orientations in the second half of the data). To obtain grid-like codes for the first half of the data, we repeated the analysis but reversed the partitioning of the estimation/test data sets (i.e., we estimated grid orientations on the second data half and tested them on the first data half). Saccade-based grid-like codes were then extracted from the parametric modulation regressor (i.e., relying on the difference between each saccade’s translational direction and the mean grid orientation of the participant, whereby a smaller difference should be associated with a stronger grid-like signal within the entorhinal cortex). We then averaged the amount of grid-like coding of all saccades within a trial, producing a trial-wise value for grid-like coding.

Next, to be able to perform a group-based analysis, we used the normalized, standard-space data and created a separate GLM (GLM 3). This model contained a single task regressor that captured all scene trials that were presented during the study period (modeled with a boxcar function, duration 4 s). This regressor was parametrically modulated with trial-wise grid-like coding. As above, GLM 3 included the six realignment parameters, their first derivatives, and the squared first derivatives into the design matrix. A high-pass filter with a cutoff at 128 s was applied. We then contrasted the parametric modulation regressors that captured the trial-wise fluctuations in entorhinal grid-like coding against baseline (entorhinal grid-like coding during scene > implicit baseline) and tested for group effects by submitting the individual contrast images to a one-sample *t*-test. Significance was assessed using cluster-inference with a cluster-defining threshold of *p* < 0.001 and a cluster-probability of *p* < 0.05 family-wise error (FWE) corrected for multiple comparisons. The corrected cluster size (i.e., the spatial extent of a cluster that is required in order to be labeled as significant) was calculated using the SPM extension “CorrClusTh.m” and the Newton-Raphson search method (script provided by Thomas Nichols, University of Warwick, United Kingdom, and Marko Wilke, University of Tübingen, Germany; http://www2.warwick.ac.uk/fac/sci/statistics/staff/academic-research/nichols/scripts/spm/).

### Data set obtained at the University of Vienna – the “validation study”

#### Study setup

To validate our results, we repeated the analyses in an independent data set involving different participants who took part in two separate sessions (the study was performed at the University of Vienna, Austria). First, participants completed a recognition memory task (study period and immediate test) during fMRI scanning while their eye movements were recorded. Second, their recognition memory was tested once more in the behavioral laboratory after one week (delayed test).

#### Participant sample

Fifty participants were invited to partake in the study. Four participants were excluded due to technical problems during the eye tracking recording, leaving a sample of 46 individuals. For the analysis of grid-like codes, 26 more participants were excluded due to a low number of identified saccades during the fMRI session (2 individuals < 30 detected saccades per condition) or due to signal drop-outs in the entorhinal cortex region (24 individuals < 14 voxels in ROI mask) to match the number of voxels in ROI masks of the discovery study (as we have done previously; Wagner et al., 2023), resulting in a sample of 20 participants (15 females, age range 18-29 years, mean age = 21.75 years, 18 right-handed). All participants were healthy, did not report any history of neurological and/or psychiatric disorders, had normal, or corrected-to-normal vision, and provided written informed consent prior to participation. The study was reviewed and approved by the local ethics committee of the University of Vienna (reference number 00538).

#### Recognition memory task

The recognition memory task consisted of two study-test cycles (i.e., study_1_, test_1_, study_2_, test_2_). During each study period, participants were instructed to memorize 48 scene and 48 face images (24 indoor/outdoor scenes, 24 female/male faces; scenes derived from the same stimulus set used in the discovery study above). Images were presented in the dimensions of 500 × 500 pixels on a grey background. An image was shown for 3 s, during which participants could freely view the image and was followed by a fixation cross (inter-trial-interval 2-7 s, mean 5 s). The order of images was pseudorandomized with the restriction that no more than three images of the same scene/face category were presented in succession while ensuring an equal number of scene/face images was displayed in every quartile of each study period. Across both study-test cycles, participants studied 192 images.

After each study period, participants completed a test period (i.e., the immediate test) where the 96 previously viewed (“old”) images were pseudo-randomly interleaved with 48 novel (“new”) images, with the constraint that no more than three images of the same image category (scene or face) or the same memory condition (“old” or “new”) were presented in succession. As above, each image was presented for 3 s, followed by a 4-point rating scale that prompted participants to indicate whether they recognized the image as “old” or “new”, ranging from “very sure old” to “very sure new” (duration 2 s). The next trial started after an inter-trial-interval during which a fixation cross was presented on the computer screen (duration 2-7 s, mean 5 s).

During the delayed test after one week, participants were shown all 192 images that were studied during the initial study periods (i.e., all “old” stimuli), interleaved with 96 novel, unseen images. Image presentation was again pseudo-randomized such that no more than three images of the same image category (scene or face) or memory condition (“old” or “new”) would appear in succession. In the following, we will focus on analyzing scene images only to enable a more direct comparison between discovery and validation studies.

#### Recognition memory performance (d-prime)

Depending on individual performance during the immediate or delayed recognition memory task, scene trials were marked as (1) images that were correctly judged as “old” (i.e., hits, collapsing across confidence ratings 1-2; mean ± standard error of the mean (SEM): immediate test, 86.9 ± 2.00 trials, delayed test, 84.8 ± 2.59 trials); (2) scenes that were correctly judged as “new” (i.e., correct rejections, collapsing across confidence ratings 3-4; immediate test, 9.72 ± 2.11 trials, delayed test, 11.4 ± 2.62 trials); (3) scenes that were mistakenly recognized as “old” (i.e., false alarms, collapsing across confidence ratings 1-2; immediate test, 3.64 ± 0.85 trials, delayed test, 5.12 ± 1.32 trials); (4) scenes that were mistakenly rejected as “new” (i.e., misses, collapsing across ratings 3-4; immediate test, 45.5 ± 0.77 trials, delayed test, 43.1 ± 1.22 trials). Individual hit and false alarm rates were *z*-scored separately for the immediate and delayed test. To avoid an indeterminate *d’* (this problem arises with hit or false alarm rates of 0 or 1), we used the log-linear approach, where 0.5 is added to both the number of hits and false alarms, while 1 is added to the number of signal and noise trials, before calculations are performed (Stanislaw & Todorov, 1999). To compensate for an unequal number of signal and noise trials in this study’s design (signal trials = 96 old images, noise trials = 48 new images at test), we added proportional values to the number of hits and false alarms (i.e., 0.7, to the number of hits and 0.3 to the number of false alarms; 2 × 0.7 to the number of signal trials, 2 × 0.3 to the number of noise trials). Recognition memory performance was then quantified using *d*-prime, calculated as the difference between these adjusted hit and false alarm rates [*z*(hits) – *z*(false alarms)]. Analysis of recognition memory performance (*d*-prime) was carried out in MATLAB (The Mathworks, Natick, MA, USA, R2020b, dprime_simple.m, version 1.1.0.0 by Karin Cox) and R (versions, base, version 4.3.0, dplyr, version 1.1.3).

#### Eye tracking data acquisition, analysis, and saccade detection

Saccades were tracked by recording horizontal and vertical eye gaze and pupil size with a video-based infrared eye tracker (EyeLink 1000 Plus, SR Research, Ontario, Canada). To map raw eye movement data onto screen coordinates, we implemented a calibration-validation procedure (as described for the discovery study). Eye tracking data were processed using Fieldtrip (https://www.fieldtriptoolbox.org).

Saccades were excluded if they were preceded or followed either by blinks (± 300 ms) or other saccades (± 100 ms). Blinks were defined as pupil dilations deviating more than one standard deviation from the mean pupil diameter. We counted a total of 19818 saccades in the eye tracking data (*N* = 46; average number of saccades per participant, mean ± SEM: 430.8 ± 15.03 saccades).

#### MRI data acquisition

Imaging data were collected at the Neuroimaging Center of the University of Vienna, using a 3T Skyra MR-Scanner (Siemens, Erlangen, Germany) equipped with a 32-channel head coil. On average, we acquired 396.77 (± 7.24 SD) T2^*^-weighted blood oxygenation level-dependent (BOLD) images during each of the two study periods and 732.54 (± 9.10 SD) BOLD images during the two immediate test periods of the recognition memory task. We used the following partial-volume echo-planar imaging (EPI) sequence: TR = 2029 ms; TE = 30 ms; number of slices = 30 axial slices; slice order = interleaved acquisition; FoV = 216 mm; flip angle = 90°; slice thickness = 3 mm; in-plane resolution = 2 × 2 mm, using parallel imaging with GRAPPA acceleration factor of 2. Slice orientation was parallel to the line connecting the anterior and posterior commissure (AC-PC alignment), with a 10° rotational shift upwards. The T1-weighted structural image was acquired using a standard magnetization-prepared rapid gradient-echo (MPRAGE) sequence with the following parameters: TR = 2300 ms; TE = 2.43 ms; FoV = 240 mm; flip angle = 8°; voxel size = 0.8 mm isotropic. We additionally acquired a T2-weighted structural image used to delineate the entorhinal cortex. A turbo-spin-echo (TSE) Sampling Perfection with Application optimized Contrasts was applied using different flip angle Evolution (SPACE) sequence with the following parameters: TR = 3.2 s; TE = 564 ms; FoV = 256 mm, voxel size = 0.8 mm isotropic, slices were oriented perpendicular to the long axis of the hippocampus.

Due to the local proximity to air-filled cavities, entorhinal cortices are susceptible to image distortions. To ameliorate this effect, we collected 30 images with the same functional sequence but with a reversed phase-encoding direction (thus, stretching potential image distortions into the opposite direction). Additionally, we acquired 10 whole-brain EPI images to facilitate the co-registration of anatomical EC masks to the partial-volume EPI images with the following parameters: TR = 2.832 s, TE = 30 ms, number of slices = 42 axial slices, slice order = interleaved acquisition, FoV = 216 mm, flip angle = 90°, slice thickness = 3 mm, in-place resolution = 2 × 2 mm, using parallel imaging with a GRAPPA acceleration factor of 2. As above, slices were oriented parallel to the AC-PC line with a 10° rotational shift upwards.

#### MRI data preprocessing

The fMRI data were processed using SPM12 (https://www.fil.ion.ucl.ac.uk/spm/) in combination with MATLAB (The Mathworks, Natick, MA, USA, R2020b). Structural and functional scans were manually AC-PC corrected. The first six functional volumes were then excluded to allow for T1-equilibration. The remaining volumes were slice-time-corrected to the middle slice and spatially realigned to the mean functional image (across both study-test cycles). FSL’s “topup” command (FMRIB Software Library; https://fsl.fmrib.ox.ac.uk/fsl/fslwiki/topup; Jenkinson et al., 2012) was applied to correct potential image distortions. Specifically, the mean functional image was calculated based on the 30 functional volumes (with the reversed phase-encoding direction) and was used to estimate and correct susceptibility-induced distortions. Since grid-like codes were analyzed in subject-native space, we refrained from normalizing the data. Functional images were smoothed with a 3D Gaussian kernel (5 mm FWHM).

For the whole-brain group analyses, the distortion-corrected data was normalized into standard space. The structural scan was co-registered to the mean functional image (across both study-test cycles) and was segmented into gray matter, white matter, and cerebrospinal fluid using the “New Segmentation” algorithm. All images (functional and structural) were spatially normalized to the Montreal Neurological Institute (MNI) EPI template (MNI-152) using Diffeomorphic Anatomical Registration Through Exponentiated Lie Algebra (DARTEL; Ashburner, 2007), and functional images were smoothed with a 3D Gaussian kernel (5 mm FWHM).

#### ROI definition

Left and right entorhinal cortex masks were segmented using the Automatic Segmentation of Hippocampal Subfields algorithm (Yushkevich et al., 2015; ASHS, software version 1.0.0, https://sites.google.com/site/hipposubfields/) based on each participant’s T1- and T2-weighted, high-resolution structural image. Masks were binarized and transformed into the subject-native space of the (partial-volume) functional images. To facilitate co-registration (which can be hampered by the partial-volume field-of-view), we progressed in several steps: First, participants’ T2-weighted structural scan (along with the segmented left and right entorhinal cortex masks) was co-registered to align with the mean functional image (based on the 10 whole-brain functional images we acquired). Second, the mean (whole-brain) functional image (along with the co-registered T2 image and the entorhinal cortex masks) was co-registered to the mean (partial-volume) functional image. The overlap between each entorhinal cortex mask and the corresponding (co-registered) structural and functional data was visually inspected for each participant.

Due to its location close to the lateral ventricle, the entorhinal cortex can be associated with a lower signal-to-noise ratio. To bypass this issue, only voxels that exceeded a signal-to-noise threshold of 0.8 were examined, mainly leading to the exclusion of voxels along the anterior-medial entorhinal cortex border. Participants with less than 14 voxels in the (left or right) entorhinal cortex mask were excluded from the analyses. Consequently, in alignment with the ROI from the discovery study, analyses were focused on the posterior-medial entorhinal cortex and were based on a final sample of 20 participants (mean ± SEM; left entorhinal cortex, 19.85 ± 1.09 voxels, right entorhinal cortex, 20.35 ± 1.19 voxels).

#### Analysis of grid-like codes

Grid-like codes during the study periods of the recognition memory task were analyzed identically to above. Analyses were focused on saccades during scene presentations only (saccades during face images were collapsed in a regressor-of-no-interest and were not modulated by their respective saccade direction angle). Each study period was partitioned into equal halves of estimation/test data sets and both study periods were combined into one GLM.

### Quantification and statistical analysis

Statistical analysis was carried out in MATLAB (The Mathworks, Natick, MA, USA, R2020b) and R (software version 4.3.0; https://www.r-project.org) using a set of correlations and Wilcoxon-tests (R stats 3.6.2). Unless stated otherwise, effect sizes were tested using Cohen’s *d*, and an α-level of 0.05 (two-sided) was adopted.

## Data and code availability

### Discovery study

Raw, anonymized data will be made publicly available upon completion of orthogonal projects that rely on the same data set. Until then, raw data are available upon reasonable request to the authors in accordance with the requirements of the institute, the funding body, and the institutional ethics board. All analyses are based on openly available software. Data to reproduce graphs (recognition memory task performance, eye tracking data, grid-like coding data, and whole-brain fMRI results) will be openly available at the Open Science Framework once the manuscript is accepted for publication (*links t*.*b*.*a*.).

### Validation study

Raw, anonymized fMRI and eye tracking data are available upon request to the corresponding authors. At present, participant informed consent does not allow for depositing the full data set. All analyses are based on openly available software or custom code. Source data to reproduce all graphs (behavioral performance and ROI-based results of all grid analyses) and custom code will be openly available at the Open Science Framework once the manuscript is accepted for publication (*links t*.*b*.*a*.).

## Acknowledgements

This research was funded in part by the Austrian Science Fund (FWF) [10.55776/P34775], awarded to I.C.W. For open access purposes, the author has applied a CC BY public copyright license to any author-accepted manuscript version arising from this submission. T.S. was funded by the European Union’s Horizon 2020 research and innovation programme (https://ec.europa.eu/programmes/horizon2020/; grant number 661373) and by the European Research Council (https://erc.europa.eu/, Starting Grant 802681).

## Author contributions

Conceptualization: L.P.G. and I.C.W.; Methodology: I.C.W.; Software: I.C.W.; Validation: L.P.G. and I.C.W.; Formal Analysis: L.P.G. and I.C.W.; Investigation: L.P.G., M.L., L.K., and T.S.; Resources: I.C.W., C.L., and T.S.; Data Curation: L.P.G., I.C.W., and T.S.; Writing – Original Draft: L.P.G.; Writing – Reviewing & Editing: all authors; Visualization: L.P.G.; Supervision: I.C.W. and T.S.; Project Administration: I.C.W.; Funding Acquisition: I.C.W. and T.S..

## Declaration of interests

The authors declare no competing interests.

## Notes

### Competing Interest Statement

The authors have declared no competing interest.

