## Supplementary Information for "Entorhinal grid-like codes for visual space during memory formation"

### Supplementary materials

#### Association between the total number of saccades and recognition memory performance (*d*-prime) in the validation study

In the discovery study, we found that individual recognition memory performance (*d*-prime) was coupled to the total number of saccades that participants made when viewing later remembered scene images. To test whether this was also the case in the validation study, we repeated the correlation analysis (number of saccades x *d*-prime from immediate and delayed tests, respectively). Indeed, we found a significant association between total saccade numbers and *d*-prime values from the immediate test ($N$ = 46, $r_{Pearson}$ = 0.46, 95% confidence interval (CI) = [0.20, 0.66], $p_{two\text{-}tailed}$ = 0.001; **Supplementary Fig. S1**). Total saccade numbers and *d*-prime values from the delayed test period (carried out in addition to the immediate test one week later, s. Methods) were also positively correlated ($N$ = 46, $r_{Pearson}$ = 0.50, 95% confidence interval (CI) = [0.24, 0.69], $p_{two\text{-}tailed} < 0.001$).

#### Discovery study, control analysis: No significant saccade-based grid-like codes in control regions

Across two independent studies, we found significantly increased saccade-based grid-like codes in the left entorhinal cortex as participants viewed scene images they were later asked to recognize. Even though the entorhinal cortex is known to house grid cells and most findings of grid cells or associated grid-like codes have been reported in the entorhinal cortex, there is evidence for grid-like coding in other brain regions as well. To test if our findings were specific to the entorhinal cortex, we analyzed grid-like codes in a set of control regions that are known to be involved in memory and visuo-oculomotion, including the hippocampus, the anterior thalamus, the frontal eye fields, and the visual cortex (**Methods**). We observed no significant saccade-based grid-like codes in any of these regions ($N$ = 29; hippocampus: mean ± SEM, 0.019 ± 0.028; frontal eye fields: mean ± SEM, 0.021 ± 0.038; visual cortex: mean ± SEM, 0.002 ± 0.072; Wilcoxon test, all $p_{one\text{-}tailed} > 0.05$), except for the anterior thalamus (but this result appeared driven by an outlier; $N$ = 29; anterior thalamus: mean ± SEM, 0.168 ± 0.087; **Supplementary Fig. S2**).

#### Discovery study, control analysis: Significant saccade-based grid-like codes when reversing estimation and test data sets

To show that our result was not driven by a specific data partitioning scheme, we repeated the main analysis but reversed the estimation and test data halves (i.e., we estimated grid-like codes in the second half of the data and tested grid-like codes in the first half). Results were virtually identical ($N$ = 29; bilateral entorhinal cortex: mean ± SEM, 0.087 ± 0.023, Wilcoxon test, $V$ = 340, $p_{one\text{-}tailed}$ = 0.0041, Cohen's $d$ = 0.45; **Supplementary Fig. S3A**).

#### Discovery study, control analysis: Significant saccade-based grid-like codes after controlling for differences in saccade durations

The accurate estimation of grid orientations relies on the even distribution of saccade durations along the different directions on the computer screen. As such, it is important to determine any potential biases in the distribution of saccade durations as these could greatly impact the results (that is, if a participant never made saccades in a specific direction, it is not possible to accurately estimate the grid orientation for that direction). Thus, we partitioned saccades into different directional bins with 10°-spacing and calculated the average saccade duration (in ms) for each directional bin per participant. We observed a significant bias in saccade durations in 8 out of 29 participants (i.e., a main effect of directional bin, $p < 0.05$ for 8/29). To mitigate this, we used an iterative process, and for each iteration randomly excluded 10% of the saccades until we could ensure that there was no significant difference across saccade durations between the different directional bins. Following this, we repeated the main analysis in these 8 participants (i.e., for each participant, we determined saccade-based grid-like codes using this reduced set of saccades) and performed the group analysis once more. The main result of bilateral entorhinal grid-like codes remained significant ($N = 29$, mean ± SEM, 0.109 ± 0.041, Wilcoxon test, $V = 328.5$, $p_{one\text{-}tailed} = 0.008$, $d = 0.49$, **Supplementary Fig. S3B**).

#### Discovery study, control analysis: No significant association between saccade-based grid-like codes and entorhinal BOLD activation

To check whether significant saccade-based grid-like codes were linked to an overall increase in the entorhinal BOLD signal (thus, reflecting individual differences in the signal-to-noise ratio of the fMRI data), we correlated the magnitude of grid-like codes in the bilateral entorhinal cortex with the average BOLD signal. The result indicated a negative correlation but was not significant ($N = 29$, $r_{Pearson} = -0.33$, 95% CI = [-0.62, 0.04], $p_{two\text{-}tailed} = 0.083$, **Supplementary Fig. S4**.

#### Discovery study, control analysis: No significant association between *d*-prime and saccade durations, or entorhinal BOLD activation

First, to test whether the relationship between saccade-based grid-like codes and recognition memory performance (measured as *d*-prime) was linked to specific saccade patterns (e.g., that participants with higher *d*-prime produced shorter saccades, possibly leading to lower grid-like codes), we correlated individual *d*-prime values with average individual saccade durations (in ms). This correlation was not significant ($N = 29$, $r_{Pearson} = -0.07$, 95% CI = [-0.43, 0.30], $p_{two\text{-}tailed} = 0.717$; **Supplementary Fig. S5A**). Second, to show that the brain-behavior relationship was not driven by differences in entorhinal BOLD signal (e.g., that for participants with higher *d*-prime values, the BOLD signal was lower), we correlated individual *d*-prime values with the average individual BOLD signal. The resulting correlation was also not significant ($N = 29$, $r_{Pearson} = 0.32$, 95% CI = [-0.06, 0.61], $p_{two\text{-}tailed} = 0.094$; **Supplementary Fig. S5B**).

#### Saccade-based grid-like codes and memory durability in the validation study

In the validation study, recognition memory performance was measured twice, immediately after the study period (immediate test) and one week later (delayed test). Thus, the validation data allowed us to examine recognition memory performance at both test time points (immediate and delayed *d*-prime), as well as overall memory durability. While *d*-prime values in the immediate and delayed tests were positively correlated ($N = 20$, $r_{Pearson} = 0.94$, 95% CI = [0.859, 0.978], $p_{two\text{-}tailed} < 0.0001$), there was no significant association between saccade-based grid-like codes in the left entorhinal cortex and *d*-prime in the delayed test ($N = 20$, $r_{Pearson} = -0.28$, 95% CI = [-1, 0.108], $p_{one\text{-}tailed} = 0.114$). To test whether grid-like codes were tied to memory durability, we first quantified a memory durability score (Wagner et al., 2019) that yielded the proportion of durable memories (remembered in both immediate and delayed tests) with respect to weak memories (remembered in the immediate test but forgotten in the delayed test; durable/weak). Consistent with the above, there was no significant relationship between saccade-based grid-like coding in the left entorhinal cortex and individual memory durability scores ($N$ = 20, $r_{Pearson}$ = -0.09, 95% CI = [-0.514, 0.365], $p_{two\text{-}tailed}$ = 0.698).

### Supplementary Figures

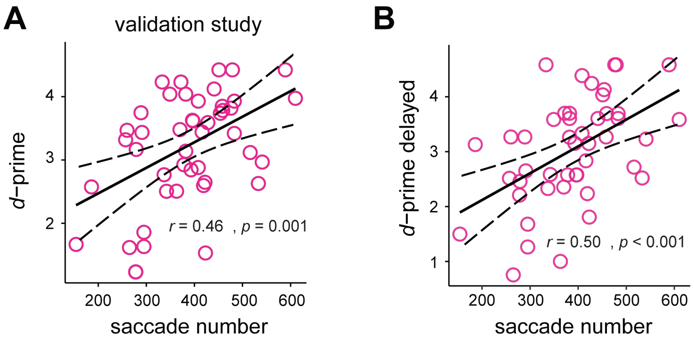


**Fig. S1: Validation study: Recognition memory performance and saccades.**
A: The scatter plot depicts the positive correlation between total saccade numbers and recognition memory performance (*d*-prime) per participant for the validation study (two-tailed, $N$ = 46, $p$ = 0.001). B: Total saccade numbers and *d*-prime from the delayed test were also positively correlated (two-tailed, $N$ = 46, $p < 0.001$). Confidence intervals (CI 95%) are indicated by the dashed line.

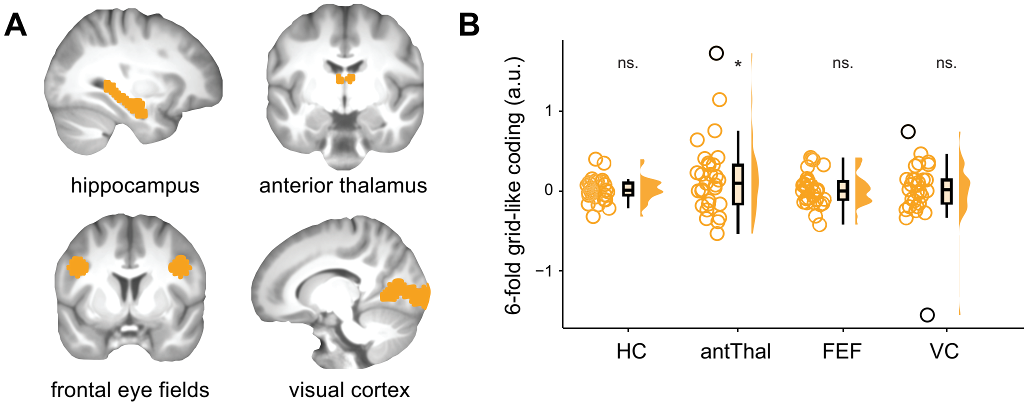


**Fig. S2: Discovery study: No significant saccade-based grid-like codes in control regions.**
A: To determine whether saccade-based grid-like codes were significantly increased in other regions apart from the entorhinal cortex, we defined four additional regions of interest (ROIs; projected in orange onto the normalized T1-weighted structural scan), the hippocampus (HC), the anterior thalamus (antThal), the frontal eye fields (FEF), and the visual cortex (VC). B: We did not detect significant grid-like codes in any of these control regions, except for the anterior thalamus (but this result appeared driven by an outlier; before outlier removal: one-tailed, $N$ = 29, $V$ = 295, $p$-value = 0.048, $d$ = 0.356; after outlier removal: one-tailed, $N$ = 28, $V$ = 266, $p$-value = 0.078). Data points show individual grid-like codes during study periods for each control region, and boxplots show the median (upper and lower borders mark the interquartile range, whiskers show minimum and maximum non-outlier values. Outliers (± 3 x MAD) are marked in black, *= significant at $p < 0.5$, ns. = not significant.

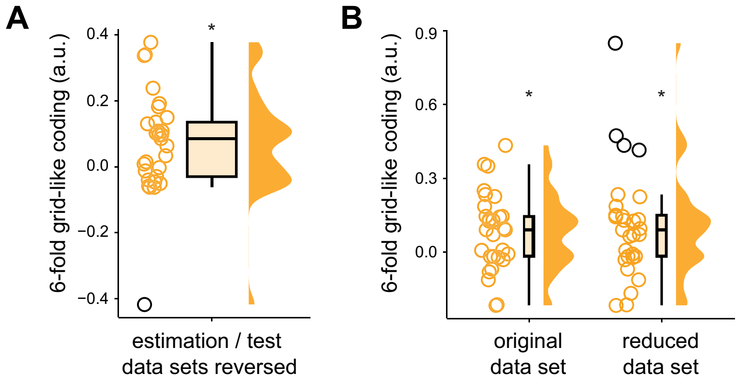


**Fig. S3: Discovery study: Significant saccade-based grid-like codes when reversing estimation and test data sets, and after controlling for differences in saccade durations.**

A: When repeating the main analysis for reversed estimation and test data halves, we again found significant saccade-based grid-like coding (one-tailed, $N = 29$, $p = 0.0041$, $d = 0.45$). B: After iteratively excluding 10% of saccades to control for individual differences in saccade durations, we observed significant grid-like coding in the bilateral entorhinal cortex (original data set: one-tailed, $N = 29$, $p = 0.006$, $d = 0.53$; reduced data set: one-tailed, $N = 29$, $p = 0.008$, $d = 0.49$). Data points show individual grid-like codes during study periods, and boxplots show the median (upper and lower borders mark the interquartile range, whiskers show minimum and maximum non-outlier values. Outliers (± 3 x MAD) are shown in black. * = significant at $p < 0.05$.

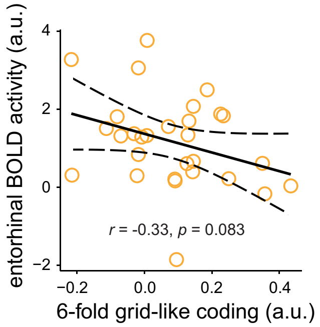


**Fig. S4: Discovery study: No significant association between saccade-based grid-like codes and entorhinal BOLD activation.**

Control analysis showing that our main result of significant saccade-based grid-like coding was not driven by changes in entorhinal BOLD signal (two-tailed, $N = 29$, $p = 0.083$). The confidence interval (95% CI) is indicated by the dashed line.

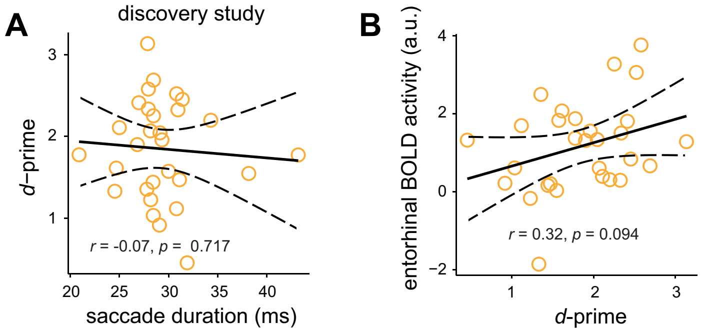


**Fig. S5: Discovery study: No significant association between *d*-prime and saccade durations, or entorhinal BOLD activation.**

A: There was no significant correlation between the average saccade durations and *d*-prime values across participants in the discovery study (two-tailed, $N$ = 29, $p$ = 0.717). B: Higher *d*-prime values were not driven by variations in the overall BOLD signal across participants (two-tailed, $N$ = 29, $p$ = 0.094). Confidence intervals (95% CI) are indicated by the dashed line.
